# An Update on Active and Passive Surveillance for African Swine Fever in the Dominican Republic

**DOI:** 10.1101/2024.10.16.618713

**Authors:** Rachel A. Schambow, Nianet Carrasquillo, Silvia Kreindel, Andres M. Perez

## Abstract

African Swine Fever (ASF) is a viral, hemorrhagic disease of swine that is reportable to the World Organisation for Animal Health. Since 2007, ASF has been expanding globally and caused severe disruption to the global swine industry. In 2021, ASF was detected in the Dominican Republic, prompting an emergency response from local and international officials. Nearly three years later, ASF is still present in the country despite control efforts. This study used data from January 2023-March 2024 from government-mandated sampling of commercial farms, a newly initiated active surveillance program of backyard farms, and passive reports to provide a comprehensive and descriptive assessment of ASF in the Dominican Republic. The attack rate for each region was calculated, and the reproductive ratio (R_0_) was estimated using the doubling time method. ASF continues to be distributed throughout the Dominican Republic with lower attack rates in central regions and R_0_ nearing 1. These results suggest that ASF in the country is reaching a stable state that does not resemble an epidemic situation. This may suggest the need to change from an approach of emergency response to one of sustained and progressive control, ultimately for the long-term goal of ASF eradication in the Dominican Republic.

## 1. Introduction

African Swine Fever (ASF) is a hemorrhagic disease of swine caused by the ASF virus (ASFV), a double-stranded DNA arbovirus of the family *Asfarviridae* ^1,2^. The ASFV only infects swine, including domestic pigs, wild boar, warthogs, bushpigs, and other wild swine species. ASFV is not infectious to humans and does not pose a food safety threat, but it causes high mortality in affected domestic pigs (nearly 100% with highly virulent strains). Because of its significant impacts to swine populations and severe economic consequences, ASF has been designated a reportable disease to the World Organisation for Animal Health (WOAH) ^3^. Since 2007, ASF has been rapidly spreading globally and is currently present in Africa, Europe, Asia, South East Asia, and the Dominican Republic and Haiti ^1,4^. ASF is difficult to control due to the lack of treatment or an approved and widely available vaccine, and consequently has caused severe disruption to the global swine industry in recent years ^3,4^.

In 2021, ASF was detected in the Dominican Republic (DR) and Haiti ^5-7^. Together, these countries make up the island of Hispaniola. Both countries were previously affected by ASF in 1978, which they dealt with by eradicating all susceptible swine on the island ^8^. In response to the 2021 reintroduction, the DR veterinary authority, with international support, employed a series of emergency strategies focused at containing and eradicating the disease ^5^. These control measures have included the development of laboratory capacity for ASF testing, surveillance, depopulation of infected herds, and compensation for culled animals. The ongoing outbreaks have had a significant impact on the DR’s swine industry, which is composed of over 13,000 producers that are mainly informal and backyard farms ^9^. It has been estimated that production decreased by 21% in 2022 because of ASF compared to the previous year, and that approximately $28.7 million USD was spent on indemnification to affected producers for government-mandated depopulation from June 2021 to September 2023 ^10,11^.

A recent evaluation of passive surveillance reports from the DR from November 2022-June 2023 suggested that, two years after its reintroduction, the disease is generally widespread throughout the country ^12^. Evaluation of the reproduction ratio, a metric that describes outbreak growth or decline ^13,14^, indicated that the ASF epidemic may be stabilizing in the population. However, this analysis was limited to only considering passively reported cases and does not provide information on the most recent months of the ASF outbreaks in the DR. Here, additional data from both passive reports and a recently initiated active surveillance program have been analyzed with the objective of reporting on the current status of the ASF epidemic in the DR and providing recommendations for next steps in ASF control. This work provides important information about the situation of the disease in the country for DR decision-makers in both the public and private sectors. These results will contribute to the design of prevention strategies to reduce ASF risk in the DR.

## 2. Methods

### 2.1 Data sources

Data on ASF surveillance from January 2023 to March 2024 was provided by the Ministerio de Agricultura Dirección General de Ganadería (DIGEGA) with support from the United States Department of Agriculture Animal and Plant Health Inspection Service (USDA-APHIS). The data received included the date that samples were received by the Ministerio de Agricultura Laboratorio Veterinario Central (LAVECEN), the name of the producer and/or farm name, the region and province of the farm, the type of surveillance (active or passive), and the result of real-time PCR (rt-PCR) diagnostic testing (positive or negative). These data represent official ASF testing in the Dominican Republic from January 2023 to March 2024. The swine population for each province in the DR was obtained from the 2022 national swine census conducted by the Ministerio de Agricultura and Banco Agricola with coordination and support from Organismo Internacional Regional de Sanidad Agropecuaria (OIRSA) ^9^.

Surveillance reports came from both active and passive surveillance in the DR. Active surveillance reports were from two different sources: 1) mandated regular sampling from large commercial farms (see definition below), and 2) from an ongoing active surveillance program in backyard farms with less than 25 pigs. The former source of surveillance was instituted by DIGEGA in March 2023. It requires commercial farms with 25 or more pigs and with the highest levels of biosecurity in the country to submit approved samples (whole blood, serum, spleen, lymph nodes, or kidney) to LAVECEN every 21 days, uninterruptedly, for rt-PCR testing following LAVECEN’s standard laboratory protocols ^15^. The second source of active reports, the ongoing active surveillance program, was designed and is being conducted by DIGEGA and USDA-APHIS. This program is intended to conduct surveillance in provinces where the number of samples received through passive reporting does not provide enough information to know the current status of disease in those populations. The sample size for each province and farm was estimated with a two-stage design using Cannon’s formula by assuming a farm-level design prevalence of 2% and a within-farm design prevalence of 10% with 95% confidence ^16^. All backyard farms with at least one pig were eligible for sampling. Farms were randomly selected from information collected via the 2022 census. However, at the time of field sampling, many new producers were found and sampled by the field team. Whole blood samples were collected from selected farms and submitted to LAVECEN for ASF testing via rt-PCR. The active surveillance program began in September 2023, and by the time this manuscript was written in June 2024, surveillance was still ongoing. In the data that were provided for this analysis, the active surveillance source (mandated regular sampling or from the ongoing active surveillance program) was not specified and therefore could not be analyzed here. Overall, the active reports represent sampled farms without any specific clinical signs or suspicion for ASF as a part of ongoing government-initiated surveillance. Passive surveillance was from farms that reported suspicion of ASF to the DR veterinary authority based on the presence of clinical signs and were subsequently investigated following DIGEGA’s routine protocols for sampling (collection of whole blood, serum, and tissue samples and submission to LAVECEN for rt-PCR and enzyme-linked immunosorbent assay [ELISA]-antibody testing) and verification of outbreaks.

### 2.2 Descriptive Analysis

Descriptive analysis was performed in Microsoft Excel, version 2016, and R version 4.3.1. Counts and proportions were calculated for the number of reports, including temporally by month and week (aggregated on a Monday-to-Monday basis), surveillance type (active or passive), region, province, and result. Choropleth maps depicting counts or proportions for each province were produced using ArcGIS Pro version 3.2.2. Map breaks were selected using the natural breaks (Jenks) setting. The positive rate for each week was estimated as the number of positive reports in a given Monday-to-Monday week divided by the total number of reports in the same time period.

The attack rate was estimated overall and for each region and province. The attack rate is defined as the number of new cases of a disease over the total number of units at-risk ^13^. It was calculated as the number of newly reported positive farms divided the total number of farms using the 2022 census estimates. There were five producers that had more than one positive report during the study period, and only their first occurrence of ASF was used to estimate the attack rate. There were also 15 positive cases that were missing a producer name; it was assumed that these reports were all new cases for the purpose of this analysis.

Because it was known that some farms would be sampled repeatedly during the study period (because of the requirement for samples every 21 days), the number of unique farms and the number of reports by farm were estimated. However, when samples were received by LAVECEN, farms were not consistently recorded using the same farm name or ID, and geographic coordinates were not available. Thus, without a clear way to identify repeating farms in the dataset, further analysis of the farm names was not performed and the estimated number of unique farms for each province are only approximations.

### 2.3 Estimation of reproductive ratio

The farm-level reproduction ratio (R_0_) was estimated for the country. R_0_ represents the average number of secondary infections caused by one infected unit throughout the duration of the infectiousness period and can indicate whether an epidemic is worsening, improving, or stabilizing ^17^. R_0_ > 1 indicates that each infectious unit causes more than one new infection and the outbreak is growing. R_0_ < 1 indicates that each infectious unit causes less than one new infection, and the outbreak is decreasing. R_0_ = 1 indicates an infectious unit causes only one new case, and the outbreak size does not increase or decrease which can lead to endemicity ^13^. The entire DR was considered the study area, and an infection was considered as an outbreak farm. R_0_ was estimated using all outbreaks in the current data (n=231), and an additional 96 passively reported outbreaks from November 2022-June 2023. Briefly, these data were previously provided by USDA and DIGEGA and analyzed in a prior publication ^12^. Together, these combined data allowed for estimation of R_0_ during the period from November 2022-March 2024 using 327 outbreaks.

R_0_ was estimated using the doubling time method, whereby R_0_ is calculated for each time period in which the number of cases doubled ^14,18,19^. However, R_0_ cannot be calculated for the end of the time period because the cases will not have doubled yet. Using this method, R_0_ was estimated as equation (1):

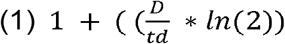

Where *t*_*d*_=the time to double the number of cases and *D*=the duration of the infectious period of the farm (days). R_0_ was estimated for each doubling interval during the study period by assuming *t*_*d*_=0 at the beginning of the time period (November 2022). Because it is difficult to estimate the duration of infection for an individual farm (as compared to an individual pig), a stochastic approach was used to account for natural variability and uncertainty in model parameters. The duration of infection was considered as the onset of infection in at least one individual pig on a farm until the depopulation of the premise. Three components were estimated: the latent (non-infectious) period of the first infected individual on the farm (Dlatent), the time for a farmer to detect and report ASF after the initial introduction (Ddetect), and the time for the veterinary authority to depopulate the farm after being notified (Ddepop). Then, D was estimated as equation (2):

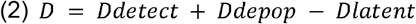

Dlatent was parameterized as Gamma(19.9, 0.3919598; mean 7.8 days) based on previous estimations of transmission parameters using herd data ^20,21^. Ddetect was parameterized as Pert(9,25,32 days), representing the minimum, most likely, and maximum time to detect ASF after introduction in a pen of 30 pigs with a total herd size of 1,200 from previous heterogenous transmission models ^22^. Finally, Ddepop was estimated as Pert(0.5,1,3 days) using expert opinion from veterinary officials in the DR with extensive, first-hand experience in responding to the outbreaks ^23^. The stochastic model was run in R version 4.3.1 using the *mcd2* package and 10,000 Markov chain Monte Carlo runs ^24^.

## 3. Results

### 3.1 Overview of Surveillance

In total, there were 9,422 surveillance reports from January 3, 2023 to March 1, 2024 (Table 1). Of these, 373 (4%) were passive reports and 9,049 (96%) were active reports. When all reports were aggregated by month, the highest number of total reports was in October 2023 (n=1,028 reports), closely followed by November 2023 (n=992) and February 2024 (n=992). Omitting March 2024 (as there are only 3 days of data), the least sampled month was April 2023 (n=448). There was also a noticeable decline in reports in the weeks of 12/26/2023 and 1/2/2024, coinciding with the Christmas and New Year’s holidays. Overall, because of the much larger number of active than passive reports, these trends closely follow changes in the number of active reports (Figure 1). The approximate number of unique farms sampled was 6,169. This number is an approximate estimation because there was no consistent farm ID and individual farms could not be easily identified or distinguished within the dataset; thus, there may be fewer unique farms than is estimated here.

**Table 1.**
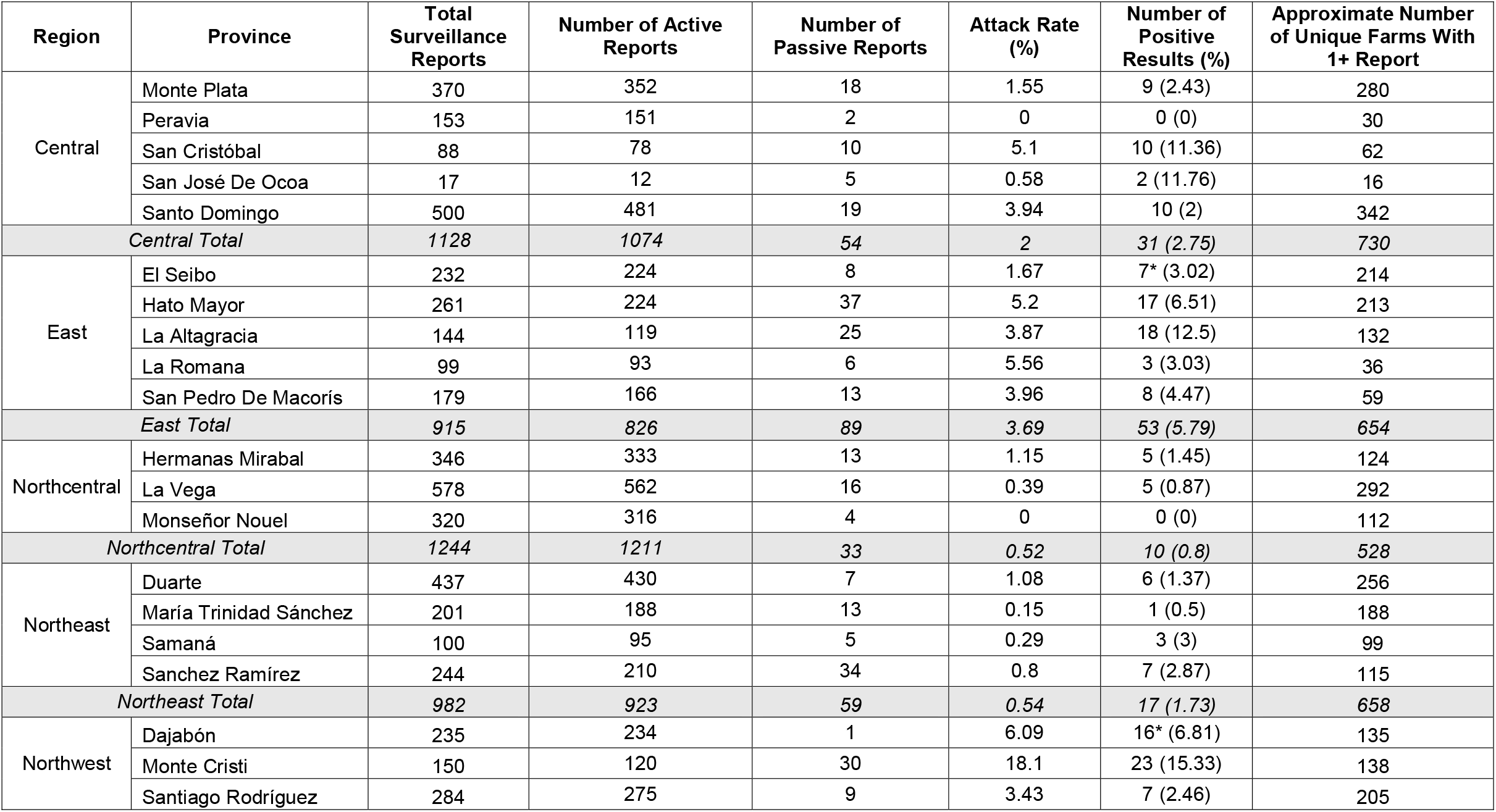
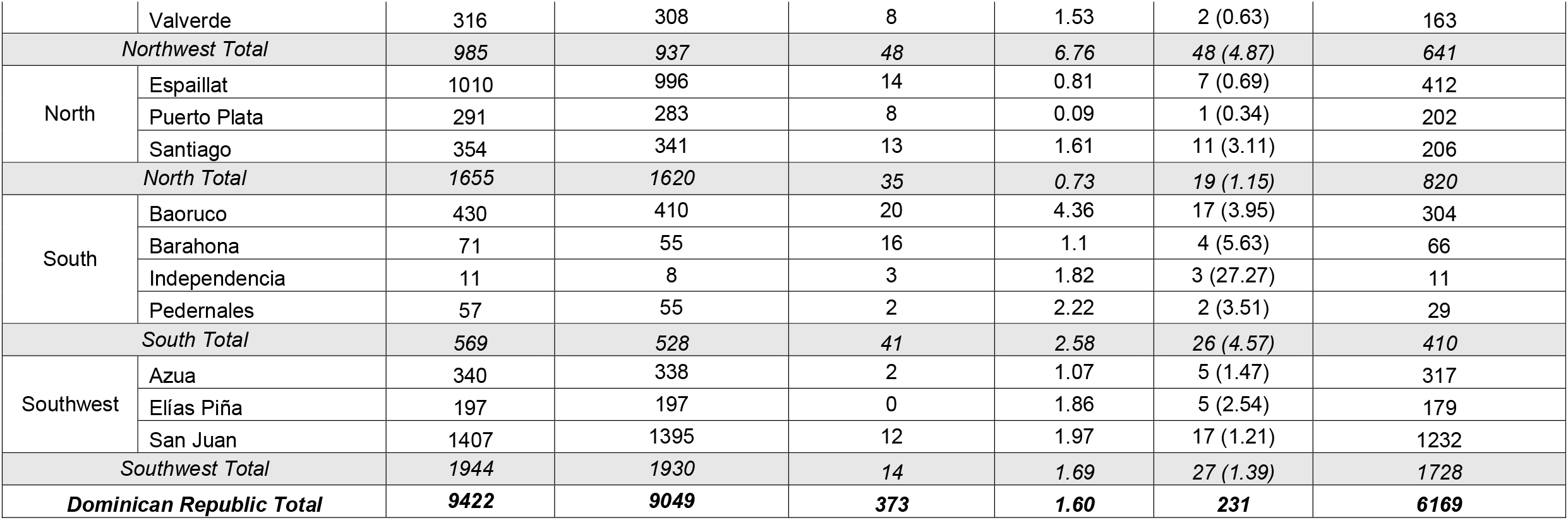
Count of African Swine Fever surveillance reports (total, active, and passive) from January 2023 to March 2024, attack rate percentage, number of positive reports and percentage of total reports, and the approximate number of unique farms that had at least one surveillance report (positive or negative), aggregated by province and region.

**Figure 1.**
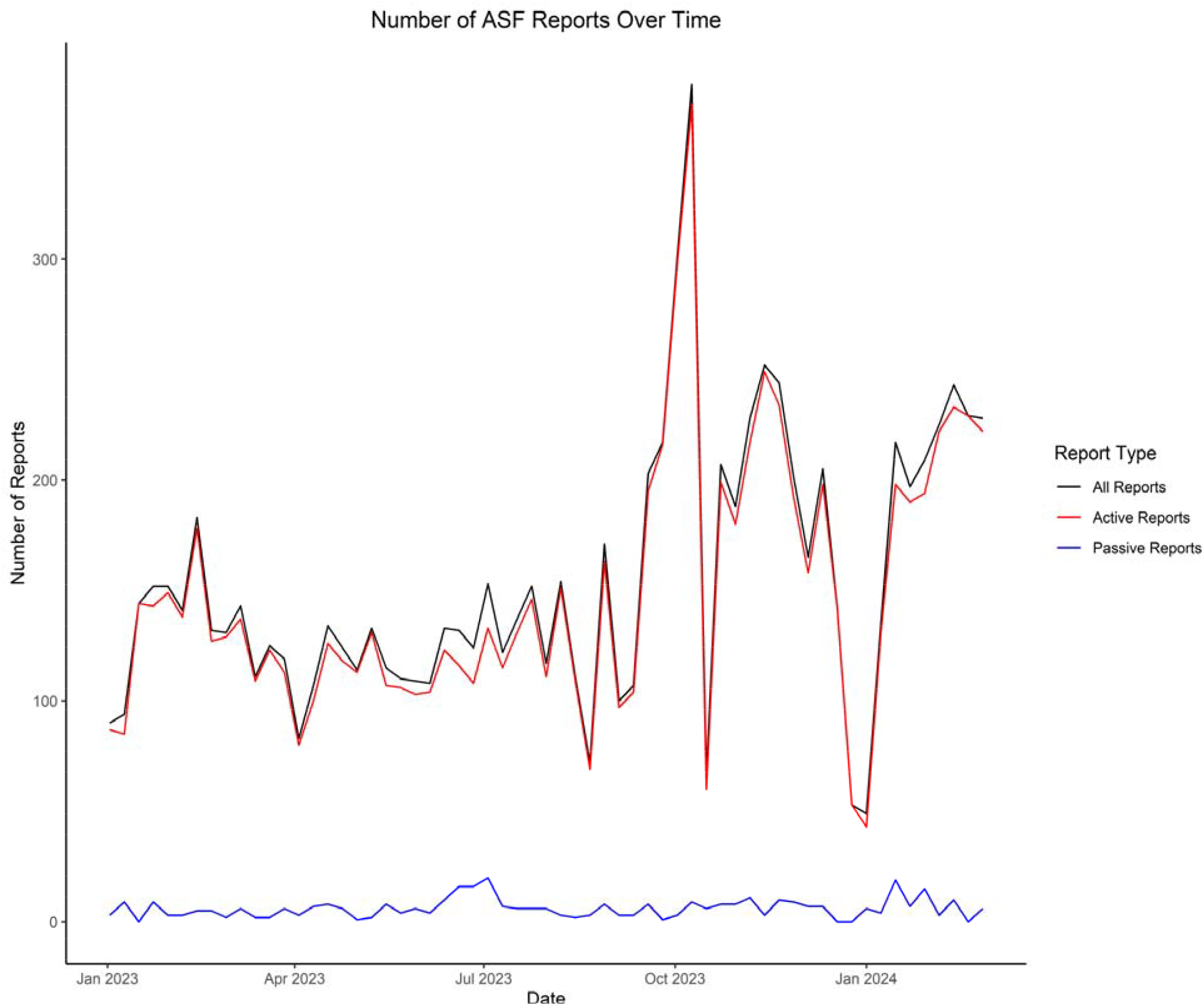
Number of African Swine Fever reports per week in the Dominican Republic from January 2023 to March 2024, considering all reports (black line), only active reports (red line), and only passive reports (blue line).

A map of provinces and administrative regions is available in Supplementary Figure 1. The Southwest region had the highest number of reports (n=1,944), whereas the South had the lowest (n=569). These trends were the same when considering the number of active reports. When considering passive surveillance only, the East region had the most passive reports (n=89) and Southwest had the least (n=14).

Not all regions were consistently reported on over time (Figure 2). The Central, East, North, Northeast, and Northwest regions had an increase in reports that coincided with the onset of the active surveillance program in September 2023. The South, Southwest, and Northcentral regions did not exhibit this same trend or not as strongly.

**Figure 2.**
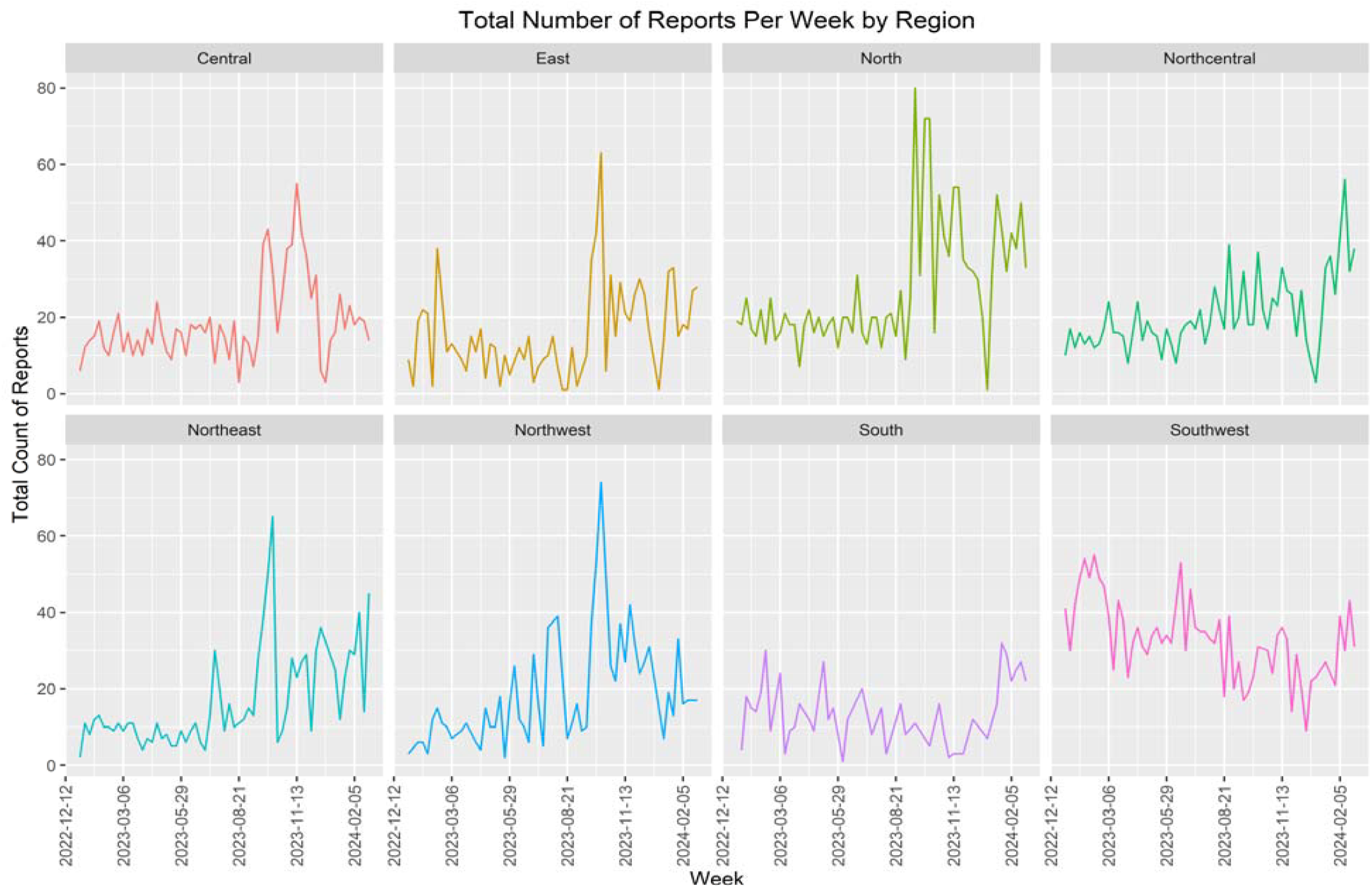
Number of African Swine Fever reports in the Dominican Republic each week from January 2023 to March 2024, by region.

### 3.2 Geographic distribution of positive results and positive rate

From January 2023 to March 2024, there were 231 positive reports (2.45%) and 9,191 negative reports (97.55%). The majority of positive reports came from passive surveillance (n=162, 70% of all positive reports; n=69 positive active surveillance reports). While it was not differentiated in the received dataset, surveillance program officials reported that all positive active reports came from backyard farms in the active surveillance program (i.e., not from commercial farms submitting samples every 21 days). There were 221 identifiably unique outbreak farms because there were five producers that appeared twice with positive results in the study period. Four of these farms had two positive reports within an approximately two-month period, and one had two positive reports within a one-week period. The highest number of positives came from the East region (n=53 total) and the lowest from Northcentral (n=10, Table 1). The provinces of Monseñor Nouel and Peravia each had no positive reports. Peravia had 153 reports from approximately 30 unique farms, and Monseñor Nouel had 320 reports from approximately 112 unique farms.

The positive rate ranged from 0.8% in the Northcentral region to 5.79% in the East region (Table 1). Across the study period, the weekly positive rate ranged from 0% (weeks of August 14-27, September 4-10, and September 25-October 1, 2023) to 8.5% (January 9-15, 2023, Figure 3). January 9-15 included 8 cases from 3 regions and 4 provinces. The three-week period from June 19-July 9 were the next highest weeks, all above 6.45% positive (n=28 positive reports).

**Figure 3.**
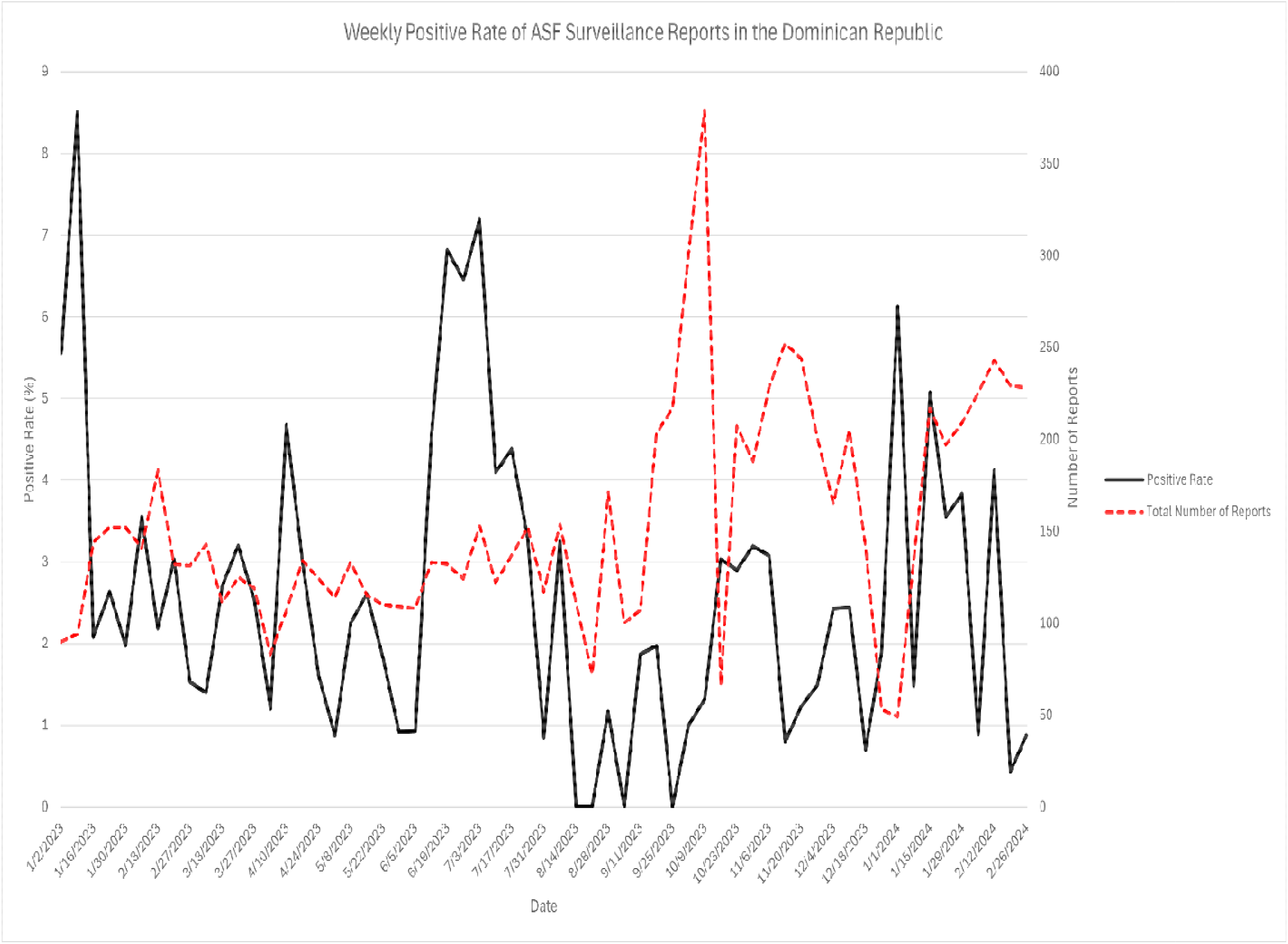
Weekly African Swine Fever positive rate of surveillance reports (black solid line) and weekly total reports (dashed red line) in the Dominican Republic from January 2023 to March 2024.

The overall attack rate for the DR was 1.60% (Table 1, Figure 4). The highest attack rates were in the Northwest (6.76%), East (3.69%), and South (2.58%) regions. The lowest were in the North (0.73%), Northeast (0.54%), and Northcentral (0.52%) regions.

**Figure 4.**
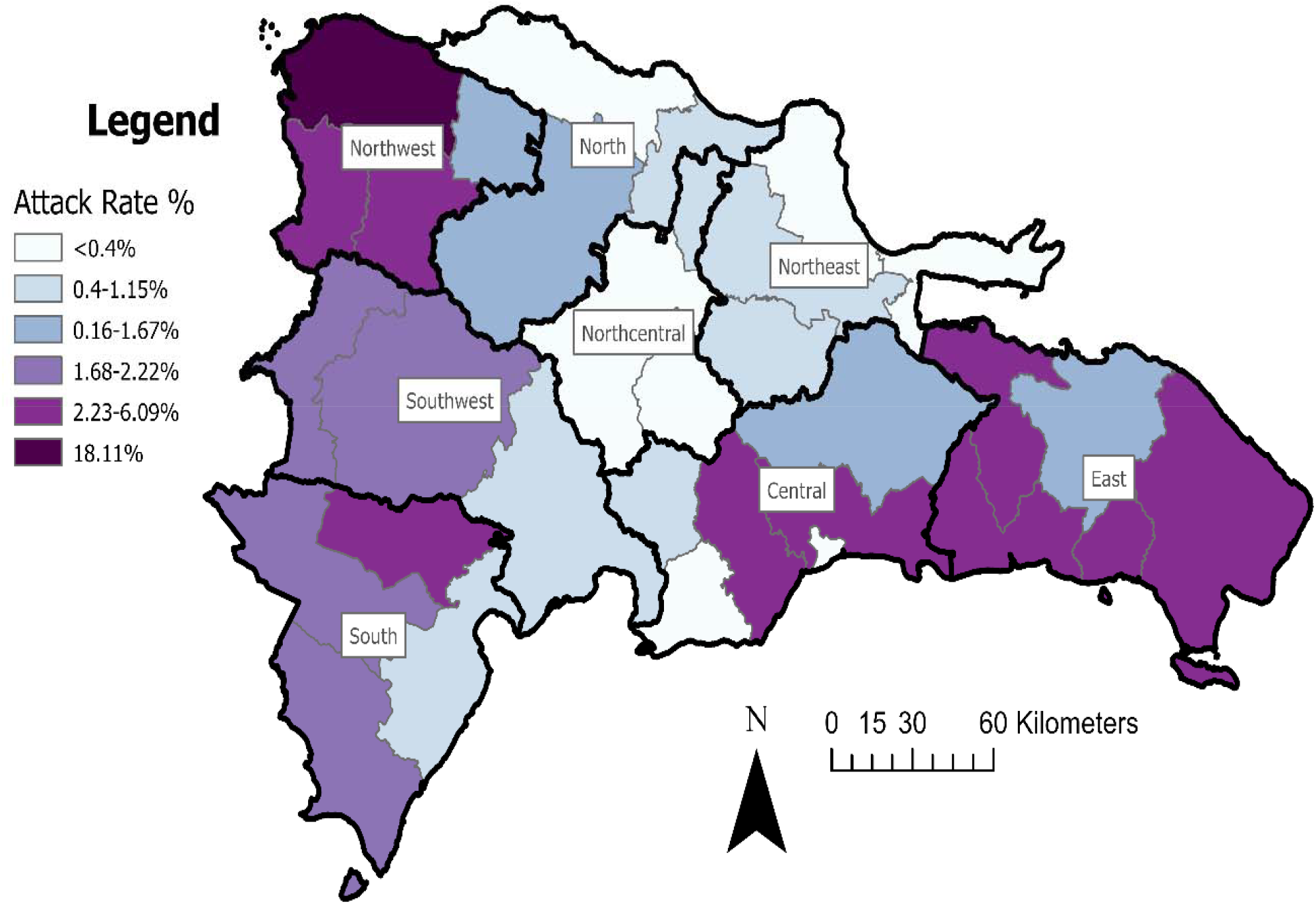
Attack rate (%) for provinces in the Dominican Republic considering outbreaks of African Swine Fever from January 2023 to March 2024. Administrative regional boundaries are designated by black, bold lines.

### 3.3 Reproduction Ratio

The median duration of infectiousness was 17.4 days (95% PI: 9.1-23.9) There were six timepoints at which the number of ASF positive reports doubled (t_d_, Figure 5). The median R_0_ estimates and their 95% PI were 4.01 (2.58-5.15), 7.03 (4.16-9.30), 1.55 (1.29-1.75), 1.37 (1.19-1.50), 1.17 (1.09-1.23), and 1.06 (1.03-1.08).

**Figure 5.**
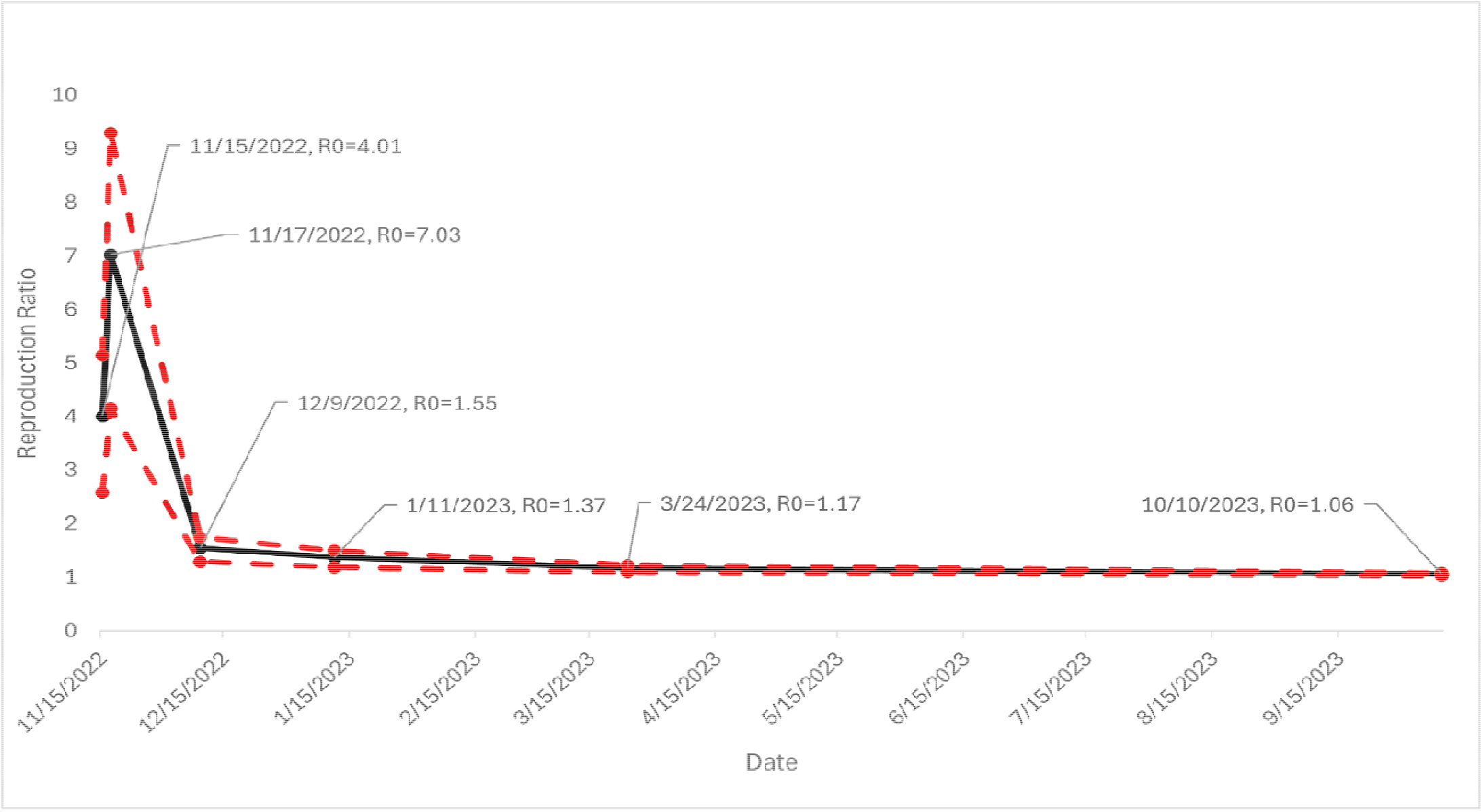
Median reproduction ratio estimates (black points) and 95% lower and upper limits of the probability interval (red points) for African Swine Fever outbreaks in the Dominican Republic from November 2022 to March 2024.

## 4. Discussion

ASF continues to be a significant disease threat globally. This update provides a timely report on the current status of ASF in the Dominican Republic that can be used by the veterinary authorities, epidemiologists, and private swine stakeholders to develop long-term ASF control programs that are tailored to the country’s current epidemiological situation.

Overall, the vast majority of positive reports came from passive surveillance. This is consistent with passive reports originating from farms with suspicious clinical signs. In 43% of passive surveillance reports, ASF was confirmed. In contrast, active surveillance focused on sampling farms without clinical signs, and only 69 (0.76%) active surveillance reports were positive. These may be farms in the earliest stages of ASF introduction and with too few clinically sick pigs to be noticed by personnel, or with infections presenting with more mild clinical signs, the latter of which has previously been reported in the DR ^5,25^. Additionally, surveillance program staff indicated these positive active reports were all from the active backyard surveillance program and not from commercial farms. It should be noted, however, that samples sent from commercial farms every 21 days are collected and submitted by producers without on-farm oversight by officials ^15^. The overall positive rate of 2.45% indicates that, compared to the initial stages of the epidemic ^26-28^, control measures have been effective at reducing prevalence and/or there has been a reduction in the susceptible population. Alternatively, some outbreaks may have been undetected because producers send pigs directly to slaughter without making a passive report.

Outbreaks occurred with relative consistency throughout the time period and were geographically widespread across administrative units (region and province). The middle part of the country consisting of the Northcentral, Northeast, and North regions had a notably lower attack rate than surrounding regions (Figure 4). According to the 2022 census, these regions contain the majority of individual producers (approximately 56% based on the 2022 census) and pigs (52%) in the country ^9^. North and Northcentral also contain over half of the country’s tecnificado or “technified” producers, those commercial farms with the highest level of biosecurity ^9^. With more commercialized producers, these regions may have more farms that implement stricter biosecurity and control measures, ultimately reducing ASF incidence. Notably, the provinces of Peravia (Central region) and Monseñor Nouel (Northcentral region) had no positive reports over the past year. It would be highly beneficial for future work to characterize the situation in these provinces and identify what swine farms may be doing there to remain ASF-free. Throughout the time period, the weekly positive rate remained under 9% without any clear trends (Figure 3). The increased positive rate in June and July 2023 included outbreaks from all eight regions and, interestingly, coincided with an increase in the total number of passive reports (Figure 1). This could suggest that there was an underlying behavioral reason or incentive for increased passive reporting of cases or could reflect changes in virus circulation.

Available data from this and a previous analysis ^12^ was also used to estimate country and farm-level R_0_ from November 2022 to March 2024 (Figure 5). R_0_ being close to one by January 2023 and remaining relatively stable through October 2023 suggests that the ASF has decreased its incidence in the country and that its impact and ecological behavior does not resemble what may be expected for an epidemic state. This is consistent with the decrease in the reported positive rate over time, from over 40% in early 2022 to 10.5% in early 2023, and then estimated here as 2.45% from January 2023-March 2024 ^26-28^. However, this decrease may be in part influenced by the large increase in sampling of farms without suspicion or clinical signs through the active surveillance program and the requirement for commercial farms to submit samples every 21 days.

There were important limitations for this work. Overall, temporal trends were difficult to assess in part because active surveillance was not consistently conducted across the country at the same time (Figure 2). Because of limited resources and personnel, geographic areas were visited at different times during the approximately 6-month period that the active surveillance program has been in progress. The lack of geographic coordinates also made spatial analysis not possible for these data. Additionally, the doubling time method used here to estimate R_0_ is traditionally better applied during the exponential or growth phase of an epidemic, whereas in this analysis, data were only available since November 2022. In reality, the outbreaks have been occurring since 2021 ^5^. Assuming no outbreaks at t_d_=0 for the purposes of the analysis may have resulted in the initial R_0_ spike which quickly returned to one afterward. Future analyses that are able to incorporate transmission modeling, geographic coordinates, or phylogenetic data may allow for a more precise estimation of R_0_ ^14,29^, which over time could indicate the efficacy of applied control measures.

Nearly three years after the reintroduction of ASF to the DR, these results may suggest the need to change from an approach of emergency response to one of sustained and progressive control. This situation draws many parallels to the ASF situation in Sardinia, where Genotype 1 virus has been present since 1978 ^30-32^. In Sardinia, ASF is thought to be maintained by complex interactions between illegal free-ranging pigs, backyard farms, and wild boar. In the DR, ASF may be similarly maintained in specific subpopulations of swine farms, such as backyard or free-ranging farms with poor biosecurity. Additionally, social conditions and limitations in the availability of human and financial resources may have prevented the full enforcement of control activities including controlling movements. This may make swift eradication highly difficult and instead may require a different approach to managing ASF in these populations, such as a combination of official registration, awareness and education, and biosecurity improvements like fencing. In a previous analysis of alternative ASF strategies in the DR, stakeholders highlighted the need to improve ASF control, such as formal registries of pig producers including for small and subsistence farmers, improved infrastructure, increased participation from the private sector, and more public-private partnerships ^33^. A progressive control plan, such as previously used for reducing foot-and-mouth disease risk in endemic countries ^34-36^, may provide a sustainable pathway to build this capacity of the DR public and private sectors and reduce ASF’s impact over time. Ultimately, this may position the DR to eventually eliminate the disease.

In summary, these results provide an evaluation of surveillance activities in the Dominican Republic from 2023-2024. ASF incidence appears to have decreased over time, though outbreaks are still being detected by both active and passive surveillance throughout the country. Management and control strategies designed for progressive control may support a long-term goal of ASF eradication in the DR.

## Acknowledgements

The authors would like to acknowledge and thank the veterinary officials and staff who conducted the surveillance and outbreak investigations and supported this data collection.

## Author Contributions

Conceptualization, S.K., and A.M.P.; Formal analysis, R.A.S. Supervision, A.M.P.; Writing—original draft, R.A.S.; Writing—review and editing, All authors. All authors have read and agreed to the published version of the manuscript.

## Data Availability Statement

Data are the property of the Dominican Republic government and have been shared with us to conduct the research here to support ASF control activities in the country. Any requests should be directed to the Dominican Republic Incident Command for ASF.

## Ethics declarations

ASF continues to be a regulated and reportable disease in the Dominican Republic. For data received for this analysis, ethical review and approval was not required because all live animal sampling procedures were performed by the Dominican Republic government (DIGEGA) as a part of its official and regular surveillance activities to control the spread of ASF. All live animal sampling was performed in accordance with international guidelines from WOAH for sampling and detection of ASF in live swine.

