## Supplementary figures and images for "An Update on Active and Passive Surveillance for African Swine Fever in the Dominican Republic"

### Supplemental File 1

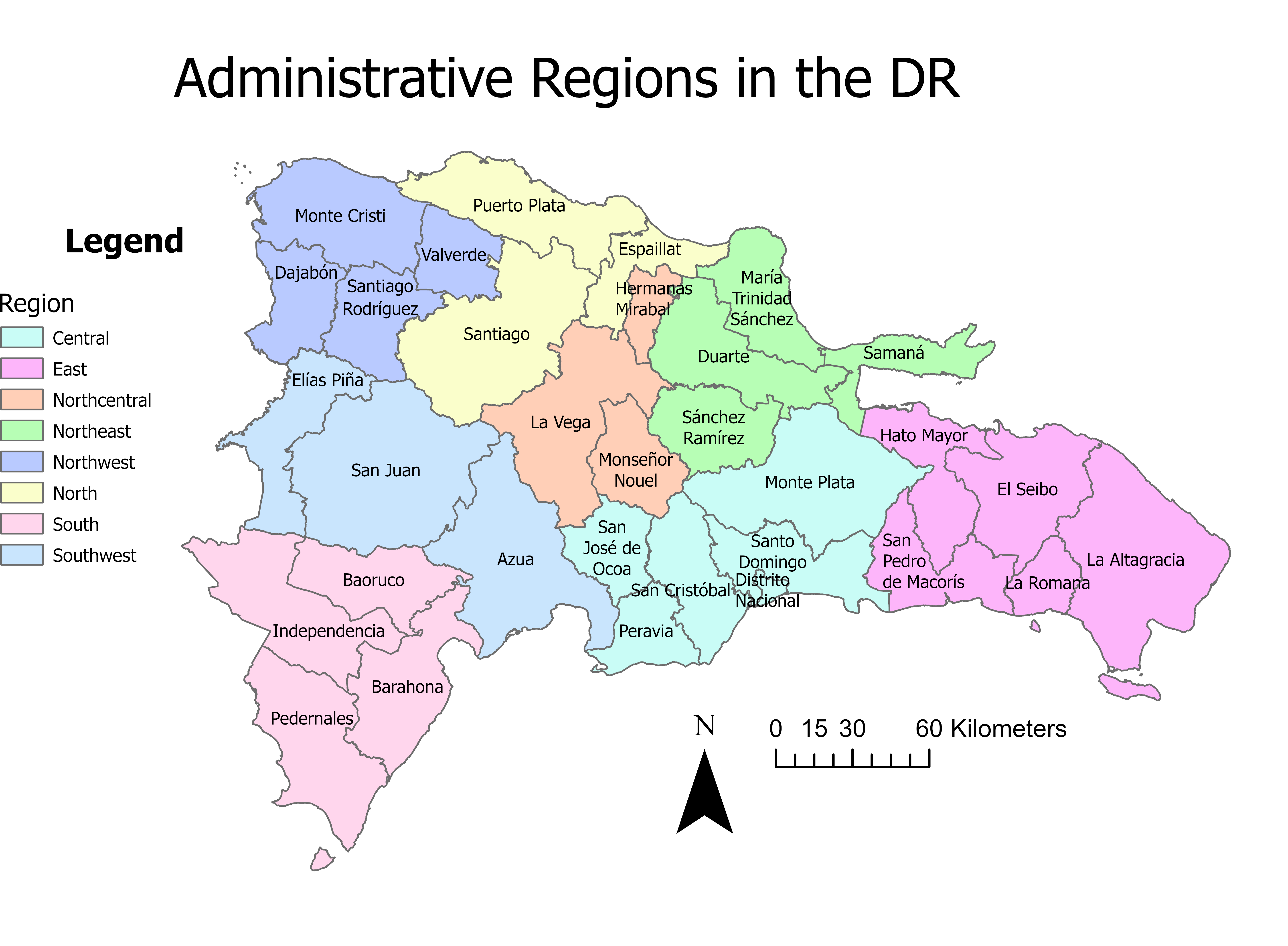
